# The Influence of Health Insurance on the Utilization of Maternal Health Care Services in Kumba Health District: A Community-based Assessment

**DOI:** 10.1101/474130

**Authors:** Mbunwe Doreen Mbuli, Mbunka Muhamed Awolu, Tanue Elvis Asangbeng, Innocent Ali Mbuli, Pepanze Jill Pangmekeh, Nsagha Dickson Shey

## Abstract

**Background:** The cash payments required to access services at health facilities called user fees are perceived to be a key barrier to improving maternal health care. This study was carried out to determine the influence of health insurance enrollment on the utilization of maternal health care services in Kumba Health District (KHD).

**Methods:** This was an analytic community-based cross-sectional study carried out in KHD, including women of child-bearing age who have had their latest delivery within 5years prior to the study. Six of the twelve health areas in the KHD were purposively selected due to the presence of health insurance companies. Probability proportionate to size sampling was used to determine the number of eligible women to be selected from each of the health areas. Data were entered into Excel 2013 and analyzed using SPSS version 20. Chi-square tests and Logistic regression were employed to identify associations between variables. Statistical significance was set at P<0.05.

**Results:** A total of 392 eligible women of child-bearing age (15-49years) were recruited into the study. A very high proportion, 94.4% of the women attended at least one ANC visit during their last pregnancy prior to the study. However, only 118(31.9%) started ANC in the critical first trimester. The prevalence of skilled facility delivery was 90.3% (95% CI: 87.2 – 93.1%). The proportion of women using a family planning method was low, 45.4% (95%CI: 49.5 – 59.2%). In multivariate analyses, insurance enrollment was significantly associated with family planning utilization (P-value: 0.000, AOR: 3.96, 95%CI: 2.14-7.35).

**Conclusion:** Less than a quarter of women of reproductive age are enrolled in health insurance in the KHD. Health insurance has been found to influence the utilization of family planning services in Kumba Health District, but not antenatal care and skilled facility delivery, though insured women are more likely to utilize these services than the uninsured. Universal health care coverage is, therefore, necessary to ensure financial accessibility to maternal health care services by all women.

## Background

The cash payments required to access services at health facilities called user fees are perceived to be a key barrier to improving maternal health care [1]. Several calls to remove the user fees have gone unfruitful in many countries[2]. Removing user fees also raises the concerns of sustainability because after removing user fees, health financing generally relies on donors or on debt relief schemes which may not be readily available[3].

Health insurance is an alternative approach to health financing, which possibly enables the removal of user fees at the point of care [4]. Unlike in high-income countries where insurance contributions are often collected through payroll deductions, this mechanism is frequently ineffective in low-income countries because a large proportion of the population is not formally employed. Instead, such contributions are increasingly raised through community-based health insurance (CBHI)[5].

This huge discrepancy in the rate of maternal deaths is due to differences in access and use of maternal health care services. For example, it is known that having a skilled attendant at every delivery can lead to marked reductions in maternal mortality and morbidity[6]. In 2010, 60% of pregnant women attended at least four antenatal care visits, 35% attended the critical first quarter visit. Underutilization of these services is primarily consequential to user fees, converging with rampant poverty in rural areas as health care financing largely depends on out-of-the-pocket payment[7]. Reducing reliance on out-of-pocket payments, by moving towards prepayment systems involving the pooling of financial risks across population groups is a significant step in ensuring financial accessibility to health care[8]. In 2006, the Cameroon government adopted a national strategy aimed at creating at least one Community-Based Health Insurance (CBHI) scheme in each health district and covering at least 40% of the population with CBHI schemes by 2015[9]. However, there is little published data on the influence of such schemes on the utilization of maternal health services in Cameroon. Findings from this study will help to inform policy makers and governments on the urgent need for universal health care coverage as a means to minimize financial inequalities in accessing maternal health care services and consequently curbing maternal mortality.

Attempts to curb maternal mortality in Cameron have paid little attention in ensuring financial accessibility to maternal health care services as a means to improve the utilization of these services. Although research has assessed the influence of health insurance on health care service utilization, little attention has been focused on its influence on the use of maternal health care services, which will be answered in this study. Findings from this study shall also highlight the urgent need for policy makers and government to implement universal health coverage as a means to address financial inequality in accessing maternal health care services. The main objective of this study was to assess the influence of health insurance enrollment on the utilization of maternal health care services among women of child-bearing age.

### Specific Objectives

1. To assess the proportion of women of child-bearing age enrolled in health insurance schemes.

2. To determine barriers to health insurance subscription among women of child-bearing age.

3. To determine the association between health insurance enrollment and the utilization of maternal health care services.

## Methodology

### Study Design and Setting

This was an analytic community-based cross-sectional study. A community based comparative assessment of health insurance enrollment and utilization of maternal health services was conducted among women of childbearing age from April 1, 2017, to August 30, 2017, in the Kumba health District.

### Study Population

Women of childbearing age (15-49 years) in the study areas were contacted and recruited into the study. Women with at least a parity of one were selected from their communities.

### Inclusion Criteria

- Included in the study, were women with at least one birth.
- They were women having any number of children with the age of the last child ranging from 0 to at most 5years.
- Those who had had a stillbirth or had lost their infant within five years were also included.

### Exclusion Criteria

- Women who were absent during the data collection period
- Women who refuse to participate

### Sample Size Determination

The sample size calculation was based on the formula of sample size determination for prevalence study[10]. Assuming Z at 95% confidence interval, and a prevalence of 0.5 **d** was assumed at 0.05 which gives us a minimum sample size of 384 participants. To take care of the anticipated non-responses of participants, 5% was added to the computed sample, given us a sample size of 404.

### Sampling Technique

Multistage sampling was employed in this study. As indicated earlier, KHD consists of 12 health areas, 06 located in Kumba city and 06 out of Kumba. The 06 health areas in Kumba city were purposively chosen because they host the three health insurance companies present in Kumba. More so, interview with the managers of these insurance companies revealed that rarely do they have clients insured from the health areas outside Kumba city. The 06 health areas consist of 11, 10, 06, 04, 12, and 09 quarters each. Probability proportionate to size sampling method was used to determine the number of participants to be selected from each health area as shown in appendix 1.

The population of women of child-bearing age in each of the health areas were gotten from the Kumba District Health Service. In each quarter, the quarter head’s house was located at a central point, and houses to the left and to the right were randomly selected. The National Immunization Days campaign numbering was used to identify houses having mothers of children 0-5 years of age, and every second house having an eligible woman was selected and the women invited into the study. If two or more eligible women were present in a household, one was randomly selected using balloting. Recruitment of study participants was done until the minimum sample size of 404 was attained. The number of participants interviewed in each quarter was proportionate to the population of the quarter. That is the quarter with high population will have a higher number of participants.

### Data Collection Tools

A structured questionnaire, modified from[11] was administered to women of child-bearing age who met the inclusion criteria and were present in the study area at the time of the study. An interview guide was used to collect information about health insurance services from the managers of the various health insurance schemes.

### Pre-testing of Questionnaire

Structured questionnaires were pre-tested two weeks before the start of the of data collection to ensure the validity of the data collection tool. Pre-testing was done among women of child-bearing age in Pulletin D, which was one of the quarters that were not selected in the study. This was to ensure that the group among whom the questionnaires were pre-tested possesses very similar characteristics to the study participants. The questionnaires were administered to 15 women of child-bearing age to ascertain its reliability.

### Data collection and processing

#### Administration of Questionnaires

A structured questionnaire was administered to consented participants. The questionnaires captured data on socio-demographic characteristics, history of utilization of maternal health care services (ANC attendance, Health Facility delivery and use of a Family Planning method) and health insurance enrollment.

#### One-on-one interview

A one-on-one interview was carried out with the managers of the various health insurance schemes to obtain information about the year of commencement of the insurance services in KHD, the MHS (mutual health services) covered by the schemes, the premiums rates, the registration and benefiting procedures

### Study Procedures

#### Fieldwork

Eligible participants gave their written consent by signing consent forms before being interviewed, and for participants less than 21 years old, assent was granted from husbands, parents or guardian after the purpose and significance of the study were explained to them. The interview was done in Pidgin English.

Information was obtained from eligible participants with the use of semi-structured questionnaires which were interview-administered. It took about 10-15 minutes to get responses. Doubts were clarified accordingly during the administration of the questionnaires

### Data Management and Statistical Analysis

#### Data Management

##### Questionnaire Handling and Storage

A code was assigned to identify each participant. Research questionnaires and workbooks and other study materials were stored safely in a locker in the laboratory and secured by locking it with a lock. After collection of the data, the questionnaires and data collection forms were checked visually for completeness, obvious errors and inconsistencies were corrected. For confidentiality, the data was coded in a Laptop where only the research team had the access code.

On a weekly basis, data from completely filled questionnaires were entered into an electronic database created on Microsoft Excel 2013. Flash drives were used to back-up the data and saved data were also self-mailed. Data analysis were done using Statistical package for social sciences (SPSS) software version 20 for windows.

##### Handling of Health Insurance Records

Data collected from the insurance records of the participants were treated with strict confidentiality. These data were used solely for the purpose of the study.

#### Statistical Analysis

Continuous variables were summarized as means ± standard deviation and categorical variables were expressed as percentages using the Statistical Package for the Social Sciences (SPSS), version 20.0. Analysis of continuous variables was carried out using the procedures of descriptive statistics and later, to identify any differences the Student’s “*t*” test was used. Categorical variables were analyzed using contingency tables involving Chi-square (χ^2^) tests to identify statistical differences between the groups. Logistic regression was also employed to determine associations between predictor and outcome variables. All p-values were two-tailed, and values less than 0.05 were considered to be statistically significant. Data were presented in the form of tables and figures.

#### Ethical Considerations

Ethical Clearance was obtained from the Institutional Review Board of the Faculty of Health Sciences of the University of Buea No: 2017/028/UB/SGIRB/FHS on July 19, 2017. Administrative approval was obtained from the authority of the Faculty of Health Sciences of the University of Buea. Administrative authorization was equally obtained from the Regional Delegation of the Ministry of Public Health for the South West Region and Kumba District Health Service. The purpose of the study and the role of the participants were well explained in the consent and assent form to the participants and participation only took place after the participant had read and signed the informed consent and assent forms voluntarily. Confidentiality was ensured by using codes to identify study participants rather than their names.

## RESULTS

A total of 404 eligible women of child-bearing age (15-49 years), were recruited into the study, giving a response rate of 392 (97%). Twelve (3%) were excluded for incomplete responses.

### Socio-demographic characteristics of study participants

The age of the study participants ranged from 16 to 47 years, with a mean age of 29.16±6.32 years. The majority of participants, 211(53.8%) were of the age group 25 to 34 years. Most of the participants, were a couple, 266(67.9%) and of the Christian religion, 372 (94.9%) Secondary level of education was predominant, 165(42.7%). Majority of both the women and their partners were self-employed, 213(54.3) and 219(75.8%) respectively. Most of the study participants, 166(46.2%) and their partners, 122(55%) had monthly income in the 11,000-15,000FCFA category respectively. The majority of 304 (77.6%) of the study participants had 1 to 3 children, with the mean age of the last child being 17.5±14.9 months (Table 1).

**Table 1:**
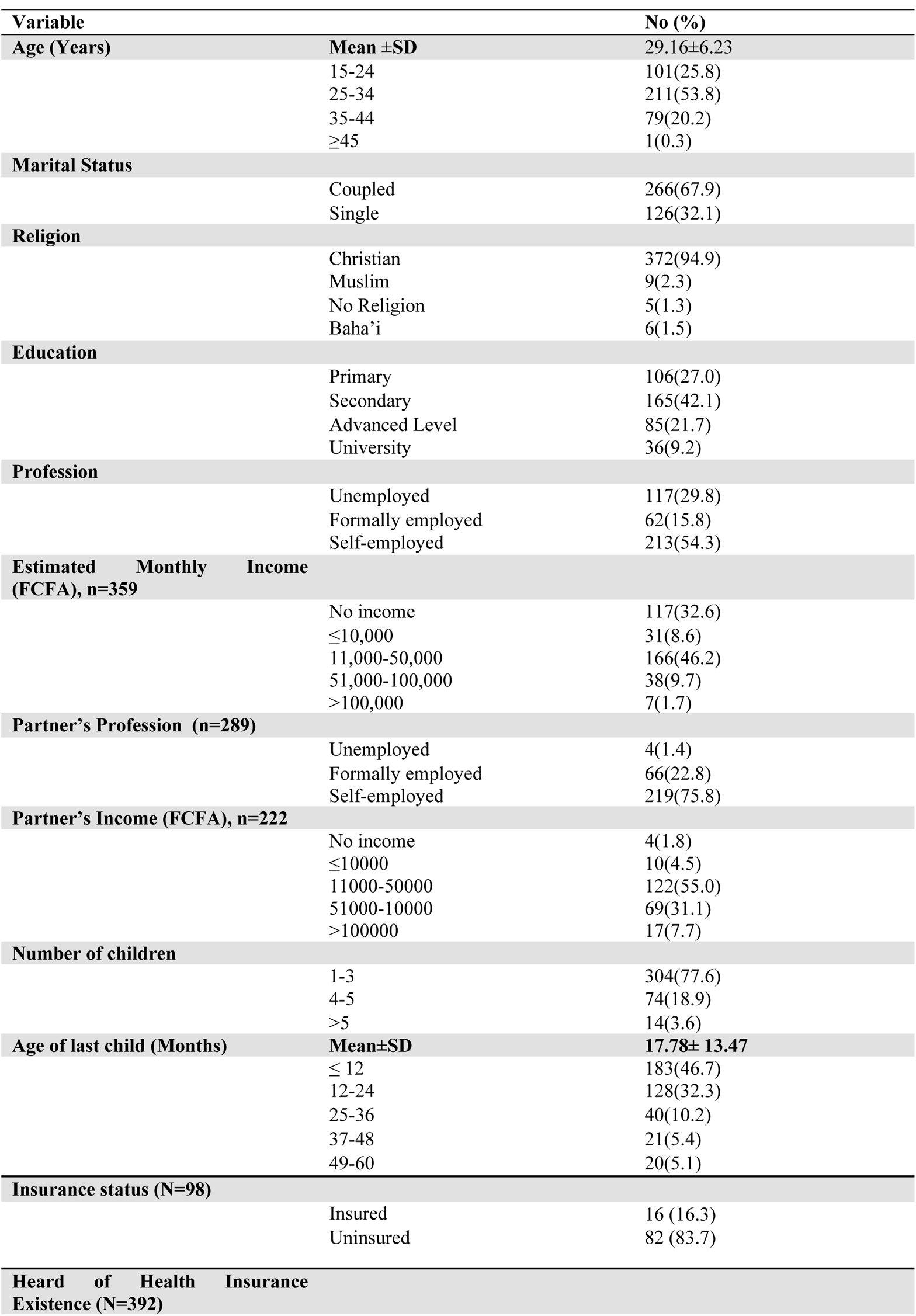
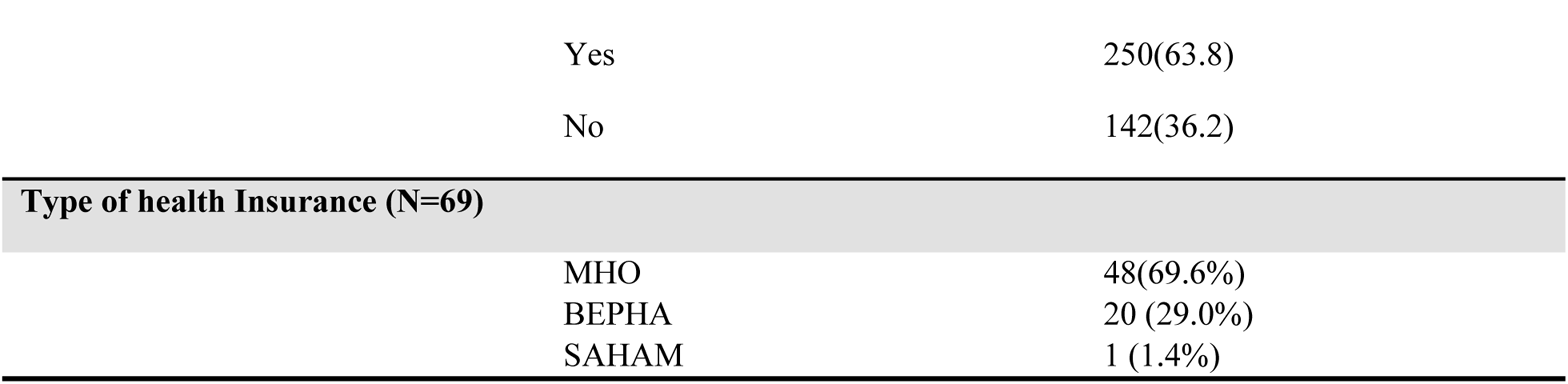
Socio-demographic characteristics of study participants (n=392)

| Variable |  | No (%) |
| --- | --- | --- |
| **Age (Years)** | **Mean ±SD** | 29.16±6.23 |
|  | 15-24 | 101(25.8) |
|  | 25-34 | 211(53.8) |
|  | 35-44 | 79(20.2) |
| | $\geq 45$ | 1(0.3) |
| **Marital Status** |  |  |
|  | Coupled | 266(67.9) |
|  | Single | 126(32.1) |
| **Religion** |  |  |
|  | Christian | 372(94.9) |
|  | Muslim | 9(2.3) |
|  | No Religion | 5(1.3) |
|  | Baha'i | 6(1.5) |
| **Education** |  |  |
|  | Primary | 106(27.0) |
|  | Secondary | 165(42.1) |
|  | Advanced Level | 85(21.7) |
|  | University | 36(9.2) |
| **Profession** |  |  |
|  | Unemployed | 117(29.8) |
|  | Formally employed | 62(15.8) |
|  | Self-employed | 213(54.3) |
| **Estimated Monthly Income (FCFA), n=359** |  |  |
|  | No income | 117(32.6) |
| | $\leq 10,000$ | 31(8.6) |
|  | 11,000-50,000 | 166(46.2) |
|  | 51,000-100,000 | 38(9.7) |
| | $> 100,000$ | 7(1.7) |
| **Partner's Profession (n=289)** |  |  |
|  | Unemployed | 4(1.4) |
|  | Formally employed | 66(22.8) |
|  | Self-employed | 219(75.8) |
| **Partner's Income (FCFA), n=222** |  |  |
|  | No income | 4(1.8) |
| | $\leq 10000$ | 10(4.5) |
|  | 11000-50000 | 122(55.0) |
|  | 51000-10000 | 69(31.1) |
| | $> 100000$ | 17(7.7) |
| **Number of children** |  |  |
|  | 1-3 | 304(77.6) |
|  | 4-5 | 74(18.9) |
| | $> 5$ | 14(3.6) |
| **Age of last child (Months)** | **Mean±SD** | **17.78± 13.47** |
| | $\leq 12$ | 183(46.7) |
|  | 12-24 | 128(32.3) |
|  | 25-36 | 40(10.2) |
|  | 37-48 | 21(5.4) |
|  | 49-60 | 20(5.1) |
| **Insurance status (N=98)** |  |  |
|  | Insured | 16 (16.3) |
|  | Uninsured | 82 (83.7) |
| **Heard of Health Insurance Existence (N=392)** |  |  |

|  |  |
| --- | --- |
| Yes | 250(63.8) |
| No | 142(36.2) |
| **Type of health Insurance (N=69)** |  |
| MHO | 48(69.6%) |
| BEPHA | 20 (29.0%) |
| SAHAM | 1 (1.4%) |

The proportion of women of child-bearing age (15-49years) enrolled in health insurance (HI) schemes. The proportion of the study participants enrolled in health insurance schemes was 17.6% (table 1). One hundred and forty-two (36.2%) of the participants had never heard of the existence of health insurance (Table 1). The majority, 48(69.6%) of the participants were enrolled in a community-based health insurance scheme called Mutual Health Organization (MHO). This was closely followed by those enrolled in a faith-based insurance called Bamenda Ecclesiastical Province Health Assistance (BEPHA), 20(29.0%) see table 1.

### Perceptions and Barriers of insured participants about insurance services

Table two describes the study participants’ perceptions of the services rendered by the various health insurance schemes. Of the 69 insured participants, a great majority, 56(81.2%) admitted that the premiums were affordable to them, Ten (14.5%) did not know which maternal health services were being covered by their health insurance. More than three quarters, 54 (78.3%) attested that insurance had made maternal health services more affordable to them. Less than half, 24(34.8%) of the insured women declared that they were not satisfied with the health insurance services, the most common reason, 13(54.2%) for dissatisfaction being complicated benefiting procedures.

The main barrier, 200(61.9%) to health insurance enrollment stated by the study participants was lack of information about the existence and importance of health insurance schemes. This was closely followed by respondents who attested that insurance services were expensive, 39(12.1%). Thirty-three (10.2%) of the participants declared that the procedure to benefit from the insurance schemes was complicated (Table 2).

**Table 2:** Participants’ perceptions and barriers to health insurance services.

| Variable |  | No (%) n=69 |
| --- | --- | --- |
| Premium Affordable | Yes | 56(81.2) |
|  | No | 13(18.8) |
|  | **Total** | **69(100)** |
| MHS Coverage | First ANC visit only | 3(4.3) |
|  | First ANC and delivery | 20(29.0) |
|  | All ANC and Delivery | 10(14.5) |
|  | Delivery | 26(37.7) |
|  | Don't Know | 10(14.5) |
|  | **Total** | **69(100)** |
| HI made MHS affordable | Yes | 54(78.3) |
|  | No | 15(21.7) |
|  | **Total** | **69(100)** |
| Satisfied with HI Services | Yes | 45(65.2) |
|  | No | 24(34.8) |
|  | **Total** | **69(100)** |
| Barrier to insurance | Expensive | 39(12.1) |
| **enrollment** | Complicated benefiting procedure | 33(10.2) |
|  | High premium | 14(4.3) |
|  | Not necessary | 34(10.5) |
|  | Not informed | 200(61.9) |
|  | Complicated registration procedure | 3(0.9) |
| **Total** |  | **323(100)** |

### Strategies to encourage health insurance enrollment as perceived by study participants

A great majority 258 (83.4%) of the women affirmed that sensitization was the way forward to enhancing health insurance enrollment. Sixty-three (19.7%) said premiums should be reduced, while 37 (11.4%) stated that the procedure to benefit should be made easy (Figure 1).

**Figure 1:** Participants’ perceptions of strategies to improve insurance enrollment.

### The relationship between socio-demographic characteristics and health insurance enrollment

The age group 25-34 was significantly associated with insurance enrollment (p=0.007) Employment, income and marital status were also positively associated with health insurance enrollment among the study participants, P=0.000, P=0.000 and P=0.033 respectively. Educational attainment, partner’s income and family size did not have any association with insurance coverage (Table 3).

**Table 3:** Health insurance enrollment by socio-demographic characteristics.

| Variable |  | Uninsured<br>n=323<br>No(%) | Insured<br>(n=69)<br>No(%) | Total (%) | X2 | P-value |
| --- | --- | --- | --- | --- | --- | --- |
| **Age group(years)** | Mean±SD | 28.65±6.24 | 31.52±6.20 | 30.085±6.22 | t=3.470 | p=0.001 |
|  | 15-24 | 112(28.6) | 14(3.5) | 126(32.1) | 12.159 | 0.007 |
|  | 25-34 | 163(41.6) | 34(8.7) | 197(50.3) |  |  |
|  | 35-44 | 47(11.9) | 21(5.4) | 68(17.3) |  |  |
|  | >45 | 1(0.26) | 0(0.0) | 1(0.3) |  |  |
|  | **Total** | **323(82.4)** | **69(17.6)** | **392(100)** |  |  |
| **Marital Status** | Single | 102(26.0) | 24(6.1) | 126(32.1)) | 10.502 | 0.033 |
|  | Couple | 221(56.4) | 45(11.5) | 266(67.9) |  |  |
|  | **Total** | **323(82.4)** | **69(17.6)** | **392(100)** |  |  |
| **Profession** | Unemployed | 104(26.5) | 13(3.3) | 117(29.8) | 4.845 | 0.028 |
|  | Employed | 219(55.9) | 56(14.3) | 275(70.2) |  |  |
|  | **Total** | **323(82.4)** | **69(17.6)** | **392(100)** |  |  |
| **Education** | Primary and below | 93(23.7) | 13(3.3) | 106(27.0) | 2.854 | 0.091 |
|  | Secondary and higher | 230(58.7) | 56(14.3) | 286(73.0) |  |  |
|  | **Total** | **323(82.4)** | **69(17.6)** | **392(100)** |  |  |
| **Income (FCFA)** | ≤ 50,000 | 168(69.4) | 29(11.9) | 197(81.3) | 20.042 | 0.000 |
|  | > 50,000 | 25(10.3) | 20(8.4) | 45(18.7) |  |  |

### The Association between health insurance enrollment and the utilization of maternal health care services

- **Utilization of maternal health care services by insurance status** In bivariate analysis, insured women were four times more likely to utilize ANC services compared to the uninsured, though this was not statistically significant (P value: 0.13, OR: 4.73, 95%CI: 0.63-35.76). There was also a likelihood of the insured women to commence ANC in the first trimester than the uninsured. This too was not significant (P value: 0.215, OR: 1.41, 95% CI: 0.82-2.44). Respondents who were covered by health insurance also had higher odds of attending four or more ANC visits as opposed to the uninsured, although this was equally not statistically significant (P value: 0.57, OR:1.21, 95%CI:0.63-2.30) (Table 4). Choice of skilled facility delivery was not significantly different among the insured compared to uninsured women. Nonetheless, insured women were more likely to deliver in a health facility than the uninsured. (P value: 0.451, OR: 1.46, 95% CI: 0.55-3.88). Women who had insurance coverage were more likely to perceive delivery fee as cheap as opposed to those not insured, but this was not statistically significant, (P: 0.151, OR: 1.53, 95% CI: 0.86-2.73). There was a statistically significant difference in the utilization of a family planning method between the insured and the uninsured women, (P value=0.000, OR: 4.78, 95%CI: 2.65-8.64) (Table 4).
- **Participants’ perception of maternal health service affordability by insurance status** There was a statistically significant difference in the perception of ANC affordability between insured and uninsured women. That is, insured women perceived ANC fee to be cheap as compared to the uninsured who perceived it to be expensive (P value: 0.017, OR: 0.51, 95% CI: 0.30-0.89) (Table 4)
- **Multivariate analysis of maternal health care service utilization by health insurance status** A multivariate logistic regression model was created to adjust for covariates that were identified from the associations of socio-demographic characteristics with use of maternal health services as possible confounders of the influence of health insurance on the utilization of maternal health care services. These covariates included; maternal age, income, profession, education and partner’s income. Outcome variables with a P value of < 0.2 were considered to be weakly associated with and were included in the model. Multivariate analysis reduced the likelihood of insured women attending at least one ANC visit compared to the uninsured, from four times to three times. (P value: 0.26, AOR: 3.22 95%CI: 0.42-25.0). Family planning utilization remained statistically significantly associated with health insurance enrollment (P-value: 0.000, AOR: 3.96, 95%CI: 2.14-7.35) (Table 5).

**Table 4:**
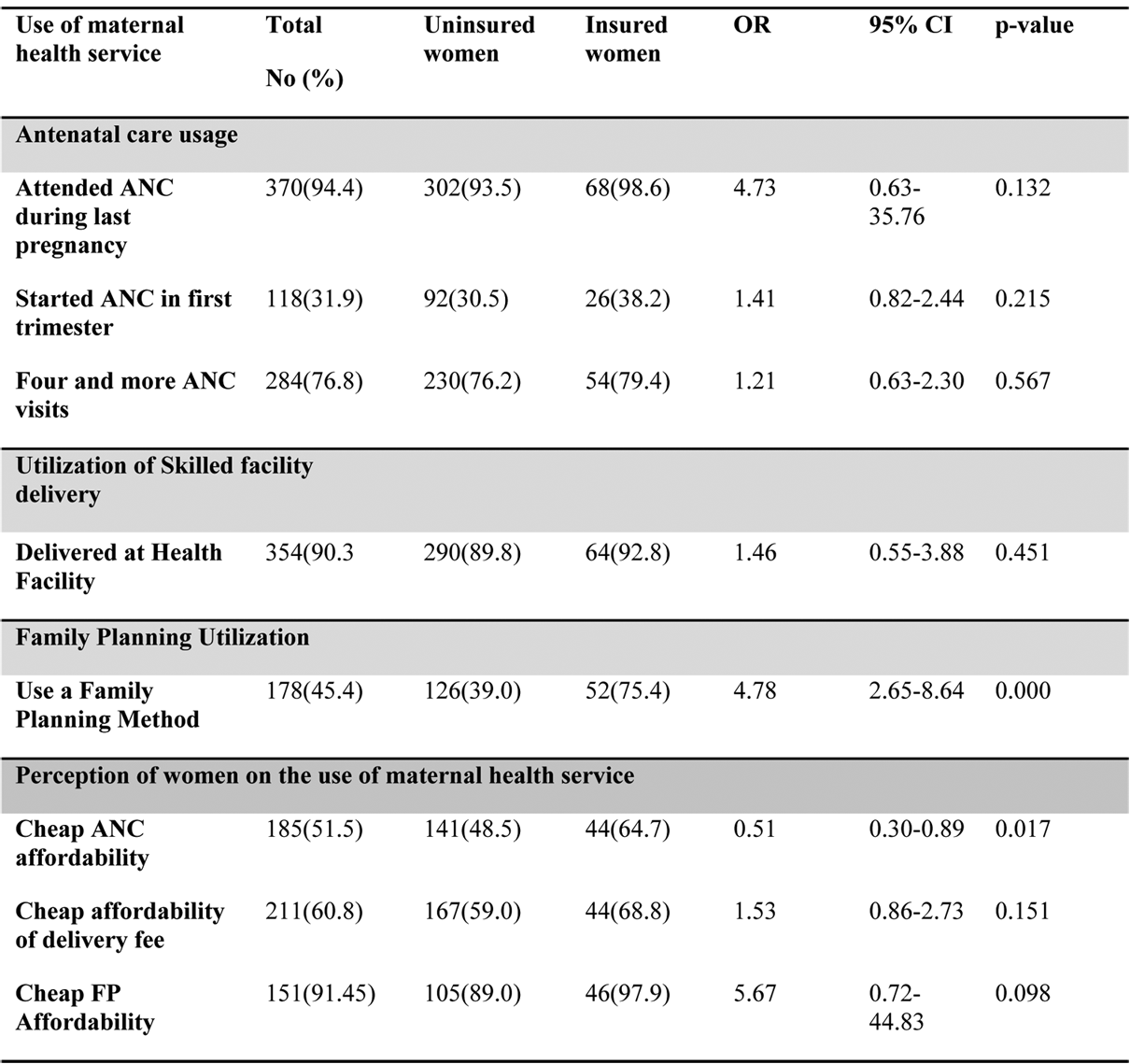
Maternal health care service utilization and perception by insurance status.

**Table 5:** Multivariate analysis of maternal health care service utilization by health insurance status.

| Use of maternal health service | Total<br>No (%) | Uninsured women | Insured women | AOR | 95% CI | p-value |
| --- | --- | --- | --- | --- | --- | --- |
| **Antenatal care usage** |  |  |  |  |  |  |
| Attended ANC during last pregnancy | 370(94.4) | 302(93.5) | 68(98.6) | 3.22 | 0.42-25.0 | 0.26 |
| **Family Planning Utilization** |  |  |  |  |  |  |
| Use a Family Planning Method | 178(45.4) | 126(39.0) | 52(75.4) | 3.96 | 2.14-7.35 | 0.000 |

**Table 6:**
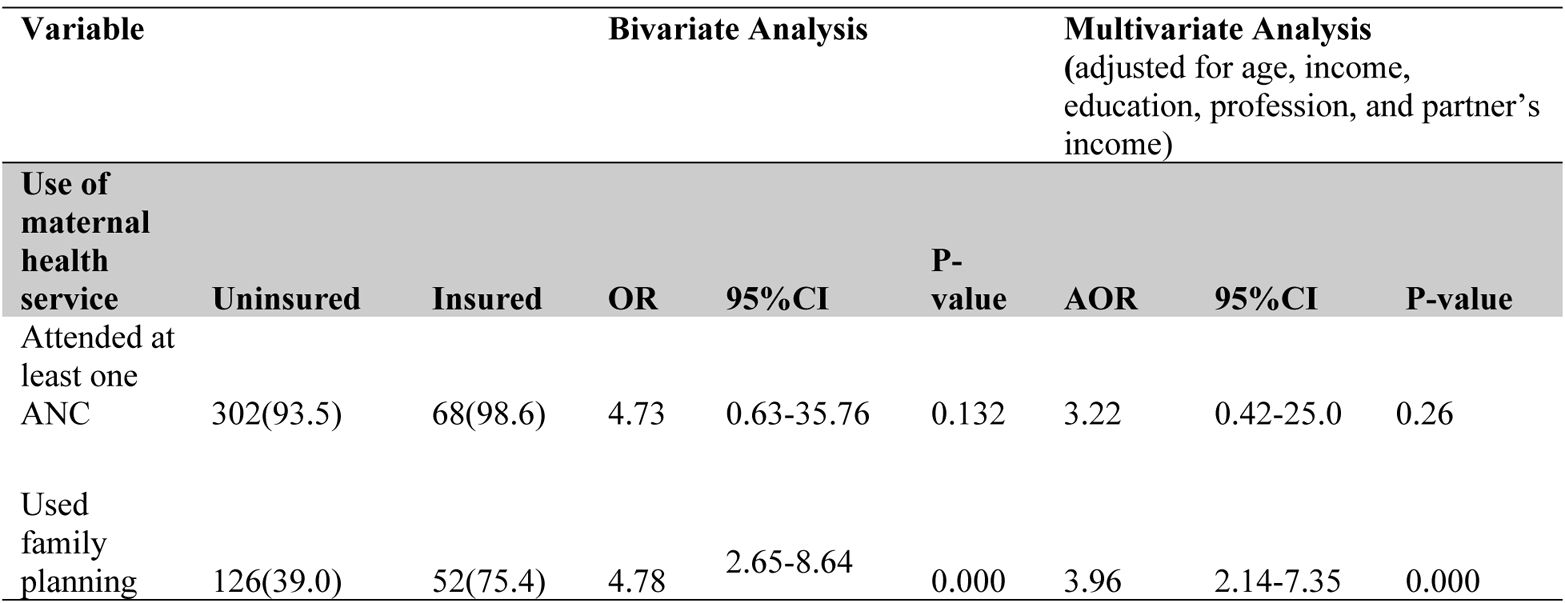
Summary of bivariate and multivariate analysis.

| Variable |  | Bivariate Analysis |  |  |  | Multivariate Analysis<br>(adjusted for age, income, education, profession, and partner's income) |  |  |
| --- | --- | --- | --- | --- | --- | --- | --- | --- |
| Use of maternal health service | Uninsured | Insured | OR | 95%CI | P-value | AOR | 95%CI | P-value |
| Attended at least one ANC | 302(93.5) | 68(98.6) | 4.73 | 0.63-35.76 | 0.132 | 3.22 | 0.42-25.0 | 0.26 |
| Used family planning | 126(39.0) | 52(75.4) | 4.78 | 2.65-8.64 | 0.000 | 3.96 | 2.14-7.35 | 0.000 |

## DISCUSSION

The main objective of this study was to estimate the influence of health insurance enrollment on the utilization of maternal health care services in Kumba Health District. Maternal health services examined were: antenatal care, skilled facility delivery and family planning. This study made use of primary data from 392 women of reproductive age (15-49 years) who had their latest delivery within 5 years prior to the study. Generally, results showed that a very low proportion of women were enrolled in health insurance schemes. A high proportion of women attended at least one ANC visit and utilized skilled facility delivery. A low proportion of study participants utilized the critical first trimester ANC visit and family planning services. Health insurance enrollment was not found to be associated with ANC and facility delivery uptake, though insured women were more likely to use ANC and facility delivery services than the uninsured. There was a significant positive association between health insurance enrollment and family planning use, confirming our research hypothesis.

To the best of our knowledge, this is one of the very few studies that have examined the influence of health insurance on ANC, Facility delivery and family planning utilization using primary data. Most studies have only assessed the association between health insurance, ANC and facility delivery, but not family planning uptake, and have made use of secondary data. The low prevalence (17.6%) of insurance enrollment in our study is an indication that health care financing in Cameroon still largely depends on out-of-the-pocket payments[7], predisposing many households to catastrophic health expenditures and propagating financial inequalities in accessing maternal health care.

In this study, four types of insurance schemes were identified in Kumba Health district: Mutual health organization (MHO), a community-based micro health insurance scheme; the Bamenda Ecclesiastical Province Health Assistance (BEPHA), a faith-based non-profit-making organization run by the Catholic denomination; SAHAM, a Moroccan private insurance company. In all the insurance schemes, premiums are fully paid for by the clients. This is contrary to what was observed in Rwanda, where the extremely poor are not required to pay premiums and in Indonesia where there is full subsidization provided for low-income households[13].

Our findings revealed that a low proportion (17.6%) of women of reproductive age is enrolled in health insurance schemes. This proportion is more than double the 7% reported in Kenya [14]. It is also far higher than the 2%, and 1% observed in Ethiopia and Burkina Faso respectively but is less than half of that obtained in Ghana and Rwanda (40% and 71%) respectively [14]. Our proportion is more than four times that reported in Cameroon (4%) in 2015[15]. This disparity could probably be due to the fact that more people are becoming informed about the need for health insurance enrollment and also based on the fact that this study was carried out in the semi-urban communities of Kumba where health insurance schemes are concentrated. The low proportion in our study could be explained by the fact that health insurance is still a new concept in Cameroon as well as many African countries and many people are not yet aware of its existence and benefits. This low insurance coverage is an that risk sharing mechanisms are poor, and many families are exposed to catastrophic health expenditures, thus aggravating the cycle of poverty in Cameroon and propagating financial inequalities in accessing maternal health care services[15].

Most of the participants, (69.6%) were enrolled in a community-based micro health insurance, MHO, followed by BEPHA, (29%). This finding is consistent with that of Wenjuan et al [13], in Rwanda, where a majority of insured women were enrolled in a community-based health insurance scheme called Mutuelle de Sante’. This could be accounted for by the fact that the CBHI schemes are run by community members who can easily be accessed, and their premiums are relatively lower. However, it contrasted that of Wang et al in Ghana, Indonesia, Albania and Cambodia [14], where enrollment into National Health Insurance Scheme was predominant. The most frequent reasons advanced for non-insurance enrollment were; lack of information on the existence and benefits of health insurance schemes (61.9%), the high cost of premiums (12.1%) and complicated benefiting procedure (10.2%). Our findings are similar to those of Jean et al [9] in Douala-Cameroon, where lack of awareness and limited knowledge on the basic concepts of CBHI, and high premiums were reasons for low insurance enrollment. Amuh A. H et al [16] in Ghana found that the major barriers to health insurance subscription included; long queues and waiting time, and negative attitude of service providers both at the healthcare facilities and the health insurance office.

Findings from this study identified the following socio-demographic factors to be associated with health insurance enrollment; age group 25-34, marital status, employment and income status had a statistically significant positive association with health insurance enrollment. This result was congruent with findings from other studies conducted in Africa[13-14], [17], where older women, those employed and of a higher household wealth were more likely to be insured than their younger, unemployed and poorer counterparts. This could be explained by the fact that poorer, unemployed women cannot afford the cost of premiums, suggesting, therefore, the need for government to get actively involved in premium subsidization and free insurance subscription for the poor and underprivileged groups. Hubert et al[16] in Ghana also found marital status to be positively associated with health insurance coverage. Contrary to our findings, Wang et al [14] in 8 African countries did not find any association between marital status and health insurance enrollment.

Insured women were three times more likely to attend at least one ANC visit compared to the uninsured, though this result did not attain statistical significance. There was also a likelihood for those insured to commence ANC in the first trimester compared to the uninsured. Those who had insurance coverage equally had a likelihood of attending four or more visits than their uninsured counterparts. These findings concur with those of Wang et al in Rwanda, Wenjuan et al in Indonesia, and Mensa et al in Ghana [4], [12], [13]. None the less, they were divergent with studies in Cambodia and Gabon, where insurance coverage had a negative impact on ANC access[14].

Our study showed no significant association between health insurance enrollment and skilled facility delivery, though insured women were more likely to utilize skilled facility delivery than the uninsured. Our findings are consistent with studies in Cambodia and Gabon, where health insurance coverage was not associated with facility delivery. However, it is divergent with findings from several other studies where there was a positive association between health insurance coverage and use of skilled facility delivery [4], [16], [18–20]. This disparity can be explained by the fact that insurance does not cover the cost of transportation to a health facility and other social costs associated with maternal health service-seeking. For example, after the user fee reform in Ghana, it was observed that many women continued to deliver at home due to the high cost of transportation to a health facility[21]. This, therefore, implies that there is a crucial need to improve geographical access to maternal health care services. The absence of an association between health insurance enrollment and facility delivery in our study could also be due to the fact that some (14.5%) of the insured women were not aware that insurance covers maternal health care services. Furthermore, insured women complained that the benefiting procedure was complicated and health facility staff preferred to attend faster to clients who did out-of-the-pocket payment than those who had to follow insurance procedures. This finding projects the need to make insurance services less bureaucratic and more electronic.

Multivariate analysis revealed that a significantly higher proportion (75.4%) of insured women used a family planning method than the uninsured (45.4%). This finding was convergent with that of Culwell et al [22], where a significantly higher proportion of insured women than the uninsured reported use of a contraceptive method (54% Vs 45%). Also consistent with our finding is that of Nearns [23] were young women with insurance coverage were more likely to use contraceptive than the uninsured. Although health insurance in this study did not cover the cost of family planning method, the association between them could be explained by the fact that insurance leads to increase access to conventional medical care, which can raise clients’ level of comfort and trust in conventional medicine [24].

## Strengths

- To the best of our knowledge, this is one of the very few studies carried out in Cameroon on the influence of health insurance on the utilization of maternal health care services.
- The study made use of primary data, therefore obtaining first-hand information on participants’ perceptions which cannot be got from secondary data which most studies used.
- The study highlights the urgent need for universal health coverage, noting that this has been declared by the director general of WHO to be the single most powerful concept that public health has to offer. Universal health coverage has been identified as a global priority in the post-2015 development agenda.

## Limitations

- The study was cross-sectional and thus inference about the causal relationship was not possible. However, it presented a snapshot of maternal health care service utilization among insured and uninsured women. Cohort studies could monitor women’s enrollment in health insurance schemes and their potential influence on maternal health service utilization.
- The presence of confounders made it challenging to attribute differences in the uptake of maternal health care services to health insurance enrollment. None the less, this was adjusted for in multivariate analysis.
- Women’s history of the utilization of maternal health care services was taken in this study hence the responses were subject to recall bias. However, a majority of the participants had their most recent deliveries below two years and could barely remember their maternal health service utilization history.
- The study was conducted only in the semi-urban communities, where health insurance services are available, hence can only give an estimate of maternal health service and health insurance coverage in the Kumba Health District. However, all six health areas in the semi-urban communities were included, which is representative of the KHD

## CONCLUSION

A very low proportion (17.6%) of women of reproductive age is enrolled in health insurance schemes, the main reason for non-enrollment being lack of information on the existence and benefits of insurance services.

Health insurance, though a new concept in Cameroon as well as other sub-Saharan African countries, has been found to influence the utilization of family planning services, but not the ANC and skilled facility delivery. However, insured women were more likely to utilize ANC and skilled facility delivery services than the uninsured. Universal health care coverage is, therefore, necessary to ensure financial accessibility to maternal health care services by all women as a means to curb maternal morbidity and mortality.

## List of Abbreviations

ANC: Antenatal care
CBHI: Community-Based health insurance
CBHI: Central Bureau of Health Intelligence
FP: Family Planning
KHD: Kumba Health District
MHCS: Maternal health Care Services
MMR: Maternal Mortality Ratio
SSA: sub-Saharan Africa
SDGs: Sustainable Development Goals
SWR: South West Region

## Declarations

### Ethical Approval and consent for participation

Ethical Clearance was obtained from the Institutional Review Board of the Faculty of Health Sciences of the University of Buea No: 2017/028/UB/SGIRB/FHS on July 19, 2017. Participants who consented for the study were interviewed. Confidentiality was respected.

### Consent for publication

Not applicable

### Availability of data and materials

The datasets used during the current study are available in the additional files section

### Author’s Contribution

MDM conceived, designed and implement the study, and partake in the data analysis, MBA draft the manuscript, NDS supervise the work. All authors. All authors contributed to the interpretation of the data and approved of the final draft.

### Author’s Information

#### Mbunwe Doreen mbuli, MPH

University of Buea, Department of Public Health and Hygiene, Buea, Cameroon, Cameroon Baptist convention health Board-Baptist convention training school for health personnel, course coordinator

Tel: (+237) 674416408

#### Mbunka Muhamed Awolu, MPH

University of Dschang, Department of Biomedical Sciences PO Box 067, Dschang, Cameroon.

Elizabeth Glaser Pediatric AIDS Foundation, Department of Research, Research Assistant. Tel: (+237) 676 534 874 / 691 730 607

#### TANUE Elvis ASANGBENG, PhD c

University of Buea, Department of Public Health and Hygiene, Buea, Cameroon Tel: (+237) 676558906, 680401388, 695953175

#### Innocent Ali Mbuli, PhD

University of Yaounde 1, The Biotechnology Center, BP 8094

#### Pepanze Jill Pangmekeh, MPH

^1^University of Dschang, Department of Biomedical Sciences PO Box 067, Dschang, Cameroon

#### Dr Nsagha Dickson Shey, Asst Prof

University of Buea, Department of Public Health and Hygiene, Buea, Cameroon,

## REFERENCES

1. Lagarde, M. and Palmer, N., “The impact of user fees on access to health services in low- and middle-income countries,” Cochrane Database of Syst Rev, no. 13(4), 2011.

2. Yates, R., “Universal health care and the removal of user fees. Lancet. 2009; 373:2078-81,” 2009.

3. Witter, S. and Adjei, S., “Start-stop funding, its causes and consequences: a case study of the delivery exemptions policy in Ghana. Int J Health Plann Manage. 2007; 22:133-43,” 2007.

4. Mensah, J., Oppomg, J. R., and Schmidth, C. M., “Ghana’s National Health Insurance Scheme in the context of the health MDGs: an empirical evaluation using propensity score matching” Health Econ, 2010.

5. Robyn, P. J., Sauerborn, R., and Barnighausen, T., “Provider payment in community-based health insurance schemes in developing countries: a systematic review,” Health Policy Plann, 2013.

6. Killewo, J. Z., Leshabari, M. T., Massawe, S. N., Jahn, A., and Mushi, D., “Use pattern of maternal health services and determinants of skilled care during delivery in Southern Tanzania: implications for achievement of MDG-5 targets.,” BioMed Central, 2007.

7. Bonono, R. C. and Ongolo-Zongo P., “Optimizing the Use of Antenatal Care Services in Cameroon. 2012. Available at: http://www.who.int/evidence/sure/FRPBCPNEN.pdf. Accessed on April 22nd, 2017” 2012.

8. WHO. Cameroon:, “Health Financing System. Geneva:” 2007.

9. Noubiap J. N., Walburga Y. A., Joel M. N, and Jean J. R, “Community-based health insurance knowledge, concern, preferences, and financial planning for health care among informal sector workers in a health district of Douala, Cameroon.,” 16:7, 2013.

10. Naing L., Winn T., and Rusli, B. N, “Practical Issues in Calculating the Sample Size for Prevalence Studies,” pp. 1: 9–14, 2006.

11. Ojong E. A., “Asseement of antenatal care service utilization in Akwaya health area: A community-based study,” Masters’ thesis submitted to the Department of Public Health and Hygiene of the Faculty of Health Sciences of the University of Buea., 2015.

12. Wang, W., Gheda, T., and Lindsay, M, “Health insurance coverage and its impact on maternal health care utilization in low and middle income countries. Demographic health survey analytical studies,” 2014.

13. Wenjuan, W., Gheda, T., and Lindsay, M., “The impact of health insurance on maternal health care utilization: evidence from Ghana, Indonesia and Rwanda,” Health Policy Plan, 2016.

14. Ramare, E., Warren, C., and Bellows, B., “Determinants of health insurance ownership among women in Kenya: evidence from the 2008-09 Kenya demographic and health survey,” Int J Equity Health., 2014.

15. Ayenika, B. M, “Primary care financing for inclusive policy reforms in Cameroon,” 2015.

16. Amuh, A.H., and Kwamena, S. N., “Health insurance subscription among women in reproductive age in Ghana: do socio-demographics matter?” among women in reproductive age in Ghana: do socio-demographics matter? PMC Health Econ Rev., 2016.

17. Mulenga, N. J., Bupe, B., and Yordanos, G., “Demographic and sicio-economic determinants of women’s health insurance coverage in Zambia,” ebph.

18. Alison, B., Lauren, A., and Laurel, E, “Effect of Health Insurance on the Use and Provision of Maternal Health Services and Maternal and Neonatal Health Outcomes: A Systematic Review,” J Health Popul Nutr, pp. 31(4):81-105, 2013.

19. Gouda, H. N., Hodge, A., Zack, W., and Jimenez-Soto, E., “The impact of health insurance on the utilization of facility-based delivery for childbirth in the Philipines,” PLoS One, 2016.

20. Yeetez, A. K., Sumiyo, O., Kwaku, P. A., Kimiyo, K., Emanuel, M., and Evelyn, A., “Factors influencing health facility delivery in predominantly rural communities across the three ecological zones in Ghana: A cross-sectional study,” PLoS One., 2016.

21. Kitui, J. G., Parker, M., Fitzpatrick, R., and Otupiri, E., “Inequities and Inaccessibility to the Utilization of Maternal Health Services in Ghana after User-Fee Exemption,” Int J Equity Health.

22. Culwel, l K. R., and Feinglass, J., “The association of health insurance with use of prescription contraceptives.,” Perspect Sex Reprod Health.

23. Nearns, J, “Health insurance coverage and prescription contraception use among young women at risk of unwanted pregnancy,” j Contraception, 2008.

24. Population Reference Bureau, “The role of health insurance in family planning,” 2014.

